# Exploring Abstract Pattern Representation in The Brain and Non-symbolic Neural Networks

**DOI:** 10.1101/2023.11.27.568877

**Authors:** Enes Avcu, David Gow

## Abstract

Human cognitive and linguistic generativity depends on the ability to identify abstract relationships between perceptually dissimilar items. Marcus et al. (1999) found that human infants can rapidly discover and generalize patterns of syllable repetition (reduplication) that depend on the abstract property of identity, but simple recurrent neural networks (SRNs) could not. They interpreted these results as evidence that purely associative neural network models provide an inadequate framework for characterizing the fundamental generativity of human cognition. Here, we present a series of deep long short-term memory (LSTM) models that identify abstract syllable repetition patterns and words based on training with cochleagrams that represent auditory stimuli. We demonstrate that models trained to identify individual syllable trigram words and models trained to identify reduplication patterns discover representations that support classification of abstract repetition patterns. Simulations examined the effects of training categories (words vs. patterns) and pretraining to identify syllables, on the development of hidden node representations that support repetition pattern discrimination. Representational similarity analyses (RSA) comparing patterns of regional brain activity based on MRI-constrained MEG/EEG data to patterns of hidden node activation elicited by the same stimuli showed a significant correlation between brain activity localized in primarily posterior temporal regions and representations discovered by the models. These results suggest that associative mechanisms operating over discoverable representations that capture abstract stimulus properties account for a critical example of human cognitive generativity.

## 1 Introduction

Generativity, the capacity to create and comprehend novel forms, is a defining feature of both language and human cognition. But what are the fundamental principles that underlie this generative behavior? Linguistic models for language processing rely on abstract linguistic variables as a means to explain this phenomenon (Chomsky, 1965). In contrast, associative models developed first in the connectionist literature (Rumelhart & McClelland, 1986) and subsequently elaborated in the deep learning (LeCun et al., 2015) and later Generative AI literatures (Kirov & Cotterell, 2018) suggest that generative behavior can emerge through the discovery of abstract features that mediate productive generalization. Both accounts propose fundamentally distinct frameworks for comprehending generativity. They diverge significantly in their interpretations of findings in linguistic, developmental, and psycholinguistic research, creating a lack of consensus on the correct paradigm (Seidenberg & Plaut, 2014). They also differ in their assertions about the nature of learning (rules or tokens), the application of this knowledge in online processing, the computations performed by brain regions (especially the left inferior frontal gyrus or LIFG), and the reliance on language-specific rules versus domain-general associative mechanisms in language processing. Both accounts offer reasonable approximations of available behavioral data because they are inherently underconstrained (Anderson, 1978), lacking decisive empirical evidence regarding the nature of neural representations and the processes they engage.

Gow et al. (2022) conducted a study to examine whether localized M/EEG data at the ROI level could be used to distinguish between abstract repetition patterns representing abstract variables or token-level abstract representations. The underlying hypothesis was that the abstracted patterns might function as linguistic variables or contribute to the representation of individual words for analogical generalization. Cluster analyses of decoding accuracy demonstrated that eight ROIs, all located in posterior temporal cortex, reliably decoded repeated syllables independently of low-level repetition activation and task demands. Further analyses indicated that the activation time series supporting decoding in various posterior MTG subdivisions causally influenced decoding accuracy in other decoder regions of STS and MTG. Importantly, these decoding processes were linked to regions associated with lexical and morphological representation (Hickok and Poeppel, 2007). However, Gow et al.’s results do not differentiate between the two accounts where activity found in the temporal areas could very well be related to the representation of variables (involved in morphology) or the representation of words; thus, the localization of decodable and causal neural information does not resolve the debate.

In this paper, we ask whether the neural abstract representations that support generativity in the Gow et al. study align with the representations discovered by a variable-free deep associative model. We will further investigate whether pretraining and task-specific performance closely parallel aspects of human neural data to test the role of associative models in simulating and comprehending cognitive generativity in human learning and representation. We ask: (i) Do variable-free network models discover the same kinds of representations that brains discover to produce the generalization of abstract syllable repetition patterns? And (ii) Is pretraining a necessary precondition for model learning

## 2 Generativity of humans and computational models

The effectiveness of any mechanistic explanation of language acquisition, use, or loss hinges on its ability to effectively tackle the issue of linguistic generativity. The robust intuitions of English speakers regarding the grammaticality of innovative, semantically challenging sentences like “Colorless green ideas sleep furiously” (Chomsky, 1957), the comparative phonological acceptability of “bnik” versus “bdik” (Chomsky and Halle, 1965), or the past tense form of the newly coined verb “wug” (Berko, 1958), all support the notion that human language is generated rather than simply memorized. However, the underlying principles governing the nature of this generative behavior are not well understood and highly debated. There are two strikingly different explanations of linguistic generativity. The Rule Account that developed in the generative linguistics tradition suggests that language users generate or model novel structures by applying language-specific abstract rules or constraints to abstract variables that capture natural classes of items (Chomsky, 1965; Jackendoff, 2002; Prince and Smolensky, 2004). Linguistic variables facilitate generalization by enabling a single computation or structural constraint to be applied to a potentially boundless range of specific instances (Jackendoff & Audring, 2020). For instance, the regular English past tense is generated by combining the variable VERB with the bound morpheme -d. This generative process does not apply to a specific verb but to the abstract variable [VERB] which can be mapped to all verbs including novel ones (Berko, 1958). In contrast, associative models developed first in the connectionist literature (Rumelhart & McClelland, 1986) and subsequently elaborated in the deep learning (LeCun et al., 2015) and later Generative AI literatures (Kirov & Cotterell, 2018) suggest that generativity is product of associative processes acting on mapping-optimized representations of individual tokens. Within this framework, the past tense of a novel form like *wug* is derived from similarity with alternations such as *walk*–*walked, talk*–*talked*, or *balk*–*balked* by characterization of discoverable/abstracted token features supporting efficient mappings.

Reduplication (the use of patterned phonological repetition to productively mark semantic and syntactic properties including intensification, plurality, and emphasis) has emerged as a core phenomena for exploring the mechanisms that support linguistic generativity (Marcus et al., 1999; Marcus, 2003; Berent et al., 2002; Berent, 2002; Rabagliati et al., 2019). It is a striking example of productivity that is widely attested in human languages (Rubino, 2013), more easily learnable than non-repetition-based forms of linguistic patterning (Berent, 2002), and most importantly, it is readily generalized to new phonological inputs that have no phonetic similarity with familiar reduplicated forms (Berent et al., 2004). Marcus et al. (1999) exposed seven-month-old infants to strings of auditory nonce words formed by repeating syllables that follow some patterns like ABB (e.g., *ga-ti-ti)* or AAB (e.g., *li-li-na*). After exposure to strings that conformed to one pattern (e.g., AAB) they used a preferential head turn paradigm to compare looking times to novel stimuli that either conformed to the exposure pattern (e.g., *wo*-*wo*-*fe*) or deviated from it (*wo*-*fe*-*fe*). Infants showed consistently longer looking times to stimuli that violated the exposure pattern, suggesting that they were able to discriminate between unfamiliar tokens on the basis of reduplication pattern. They argued that this could only be explained by rule-based processing because the lack of phonemic overlap between exposure and test items seemed to rule out similarity-based associative processes that are the primary theoretical alternative to rule-based explanations for generativity. Following Marcus’s study many studies have examined how humans discover and generalize relationships involving identity rules using artificial grammar learning paradigms (Gomez, 2002; Pena et al., 2002; Gerken, 2006; Endress et al., 2007).

To further demonstrate the necessity of rules (operations over variables), Marcus et al. (1999) also conducted simulations using a Simple Recurrent Network (SRN) (Elman, 1990) to model the generalization observed in their experiment. They noted that this variable-free model failed to replicate the infants’ behavior and concluded that this failure reflected the fundamental inadequacy of variable-free approaches to capture human (variable-dependent) processing. Subsequent attempts to model Marcus et al.’s (1999) human data using variable-free network models have met with varying degrees of success. This work has shown that model performance is influenced by various factors, including pretraining (whether the model has any prior knowledge about phonemes, syllables or any abstract relations that will help the model to figure out the task at hand) (Seidenberg & Elman, 1999a,b; Altmann, 2002), encoding assumptions (whether the model is trained on input vectors that represent phonetic features, place of articulation, vowel height, primary/secondary stress or non-featural random vectors) (Negishi, 1999; Christiansen & Curtin, 1999; Christiansen, Conway, & Curtin, 2000; Dienes, Altmann, & Gao, 1999; Altmann & Dienes, 1999; Shultz & Bale, 2001; Geiger et al., 2022), and model type (whether the model is a neural network, autoencoder trained with cascade-correlation, auto-associater, Bayesian, Echo State Network or Seq2Seq) (Shultz, 1999; Sirois, Buckingham, & Shultz, 2000; Frank and Tenenbaum, 2011; Alhama and Zuidema, 2018; Prickett et al., 2022), and task (whether the task is to predict or identify rules, words, syllables, or patterns, or segment syllable sequences into “words”) (Seidenberg & Elman, 1999a, 1999b; Christiansen & Curtin, 1999;) (see Alhama and Zuidema (2019) for a detailed review of the computational models). These factors have made it challenging to draw direct comparisons with human behavior, further fueling the ongoing discussion.

Among these factors, the role of pretraining on recurrent model acquisition of repetition-based rules deserves more discussion. Seidenberg and Elman (1999a,b) proposed that infants might have acquired the capacity to discern phonological similarity between syllables through prior exposure, and they address this by extensively pre-training an SRN with syllables, enabling the SRN to recognize identity relationship between syllables. In Altmann’s (2002) study, prior knowledge integration involved pre-training a model with 10,000 sentences from Elman (1990), wherein the model predicts the subsequent word using localist vectors, without considering syllables or phonemes. Integrating relevant prior knowledge into the initial state of the models might facilitate the learning process in converging towards the generalization that infants appear to acquire more readily. This is a valid assumption because Marcus et al.’s seven-month-old infants were not tabula rasa. Interpolating from the findings of Hart and Risley (2003), it appears that children from families on welfare are exposed to approximately 1.9 million words, children from working-class families hear about 3.8 million words, and children from professional families are exposed to approximately 6.8 million words by the age of 7 months. It is worth noting that deep learning models, driven by the principle of hierarchical feature representation, extract and organize increasingly abstract data features, similar to human cognition. This approach enhances computational efficiency and forms the foundation for pretraining, a technique where models are initially trained on a related task to learn useful features before fine-tuning the target task. However, for the validity of prior knowledge argument, it is essential to identify the precise components of prior knowledge that impact the ability to generalize to novel items. For instance, Seidenberg and Elman (1999a) incorporated pretraining into their SRN, mapping sequences of syllables to discern whether each syllable matched its predecessor. Marcus (1999) contended that this form of pretraining lacks naturalness, and Shultz and Bale (2001) emphasized that a model cannot be trained on identity relations, as it would be an unfair advantage.

It is unclear whether the limitations of existing models demonstrate the fundamental need for variables to explain this type of generativity (and by extension human performance), or whether they simply reflect the limitations of current implementations of variable -free associative models. LeCun, Bengio and Hinton (2015) demonstrated that deep learning network architectures can discover abstract features that support dramatic generativity through variable-free associative processes. While useful as a proof of concept for the potential computational adequacy of associative mechanisms to explain human generativity, questions remain about how realistic they are as neural models and as psychological models given the vast training sets, they require to achieve human-like performance. Work relating modeling to neural data has the potential to show how these computational constraints shape human neural processing. Furthermore, in the ever-evolving landscape of cognitive research, an intriguing avenue of inquiry has emerged through neural studies, delving into the intricate neural underpinnings that underlie the recognition and processing of abstract repetition patterns, adding another layer of depth to our understanding of human generativity and cognitive processes (Yang et al., 2019; Kanwisher et al., 2023).

Gow et al. (2022) provides the most direct examination of the interplay between generativity and neural mechanisms. This study tried to localize M/EEG data at the ROI level to distinguish between abstract variables vs. token-level features. A support vector machine (SVM) classifier technique that had been previously applied to MEG data was adapted to probe individual ROIs identified by Granger Causation Analysis (GCA). The analysis aimed to establish whether patterns of neural activity that could be decoded had a causal influence on downstream processes—a crucial but often overlooked criterion for determining functional roles in processing and representation (Dennett, 1987; Kriegeskorte & Diedrichsen, 2019). Data were collected during an artificial grammar learning experiment in which participants briefly encountered CV-CV-CV nonwords following a reduplication pattern (AAB, ABB, or ABA) and judged whether phonemically orthogonal nonwords followed the same rule or pattern. Behavioral results showed that participants performed the task with high accuracy. Neural analyses revealed a broadly distributed bilateral network encompassing 67 ROIs with distinct activation patterns during the task, SVMs were trained to distinguish between items based on their reduplication pattern and were subsequently tested on their ability to classify the reduplication patterns in untrained items created using different syllable sets. Cluster analyses evaluating decoding accuracy revealed that eight ROIs (see Fig. 1), situated exclusively in the posterior temporal cortex, consistently decoded repeated syllables, irrespective of low -level repetition activation and task requirements. Subsequent analyses indicated a causal relationship, demonstrating that the activation time series supporting decoding in various subdivisions influenced decoding accuracy in other regions. However, Gow et al.’s findings fail to distinguish between the two accounts, leaving open the possibility that the observed activity in the temporal areas may be connected to the representation of variables (involved in morphology) or the representation of words. Consequently, the localization of latent information does not bring resolution to the ongoing debate.

**Figure 1:**
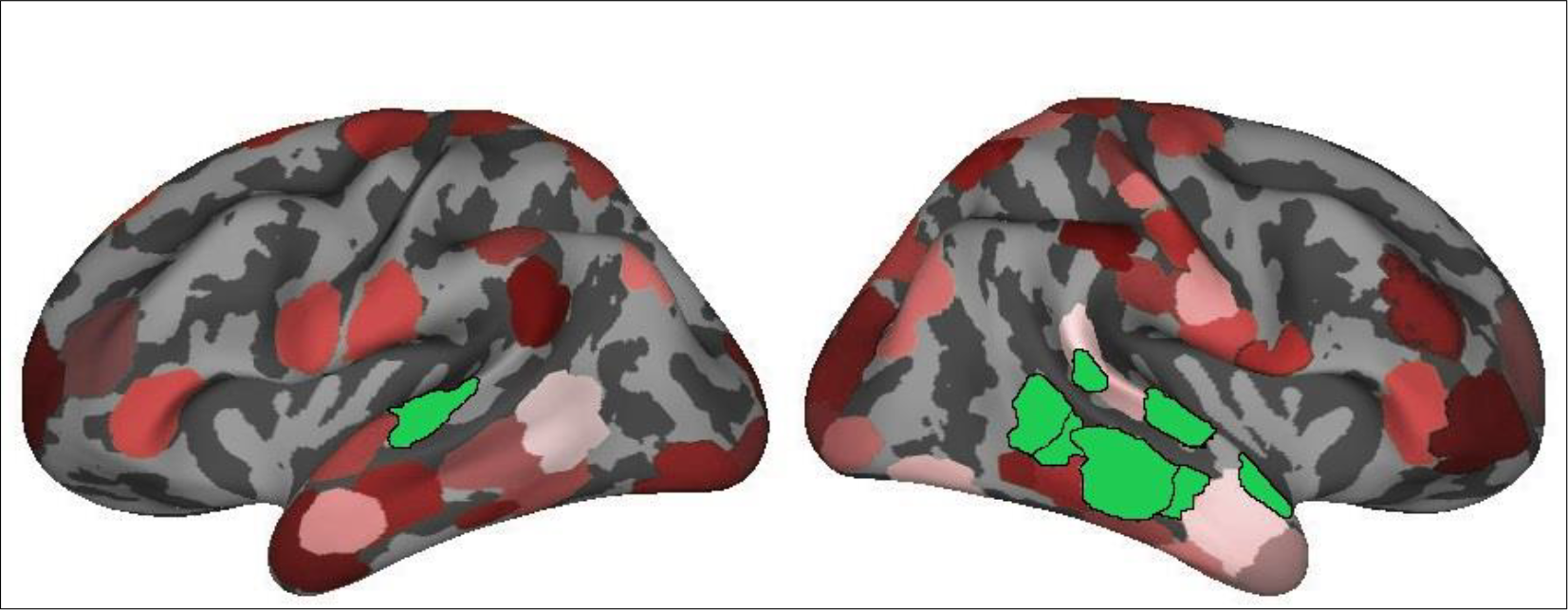
Regions of interests (ROIs), used in Gow et al. (2022), visualized over an inflated averaged cortical surface. Lateral view of the left and right hemisphere is shown. Highlighted ROIs (L_STG-1, R_STS-1 (most posterior superior), R_STG-2,3 (posterior to anterior), and R_MTG-1,2,3,4 (posterior to anterior)) showed reliable activation differences, successful decoding, or both, for reduplication.

The goal of the current study is to determine whether the abstract neural representations discovered by Gow et al. (2022) are consistent with the abstract token representations discovered by variable-free associative models. We do this by presenting a variable-free deep LSTM model trained on cochleagrams of the stimuli used by Gow et al. to discriminate stimuli based on reduplication pattern and comparing patterns of stimulus similarity within the model to patterns of ROI-level evoked activation similarity by the same stimuli in Gow et al. using Representational Similarity Analysis (RSA) (Kriesgerkorte et al., 2008; Diedrichsen and Kriegeskorte, 2017). Additionally, we explore the effects of pretraining and task-specific mapping on performance on model performance and the relationship between features discovered by the models and human neural data. To do this we trained a deep LSTM model with dropout (as explored in Geiger et al., 2022 and Prickett et al., 2022) using two distinct encoding assumptions. The first assumption involved a pattern learner trained on random vectors representing three patterns (Geiger et al., 2022). We then employed a word learner trained on vectors representing individual words based on syllable position. Consequently, we explored whether any of these variable-free network models reveal comparable representations to those identified in the brain, leading to the generalization of abstract syllable repetition patterns.

## 3 Computational Modeling Methods

Within this section, we present a detailed account of the methodological framework employed in our research, encompassing various aspects such as training data, network architecture, testing procedures, decoding techniques, representational similarity analysis, considerations of replicability, and the hardware and software infrastructure utilized for our study.

### 3.1 Training data

We used the same audio files as in Gow et al. (2022). There was a total of 23 syllables, and we used sixteen in training (/ba/, /t∫ɪ/, /dɪ/, /dʒɪ/, ka/, /nɪ/, /pɪ/, /rɪ/, /∫a/, /sɪ/, /ta/, /ðɪ/, / θu/, /va/, /zɪ/, /ʒu/) and seven in test (/fu/, /ga/, /hɪ/, /la/, /mɪ/, /wa/ and /ji/). Training data included 720 (240 for each pattern) phonemically balanced trisyllabic CV.CV.CV nonwords which were created by concatenation of sixteen different syllables following the syllable reduplication patterns: ABA (e.g., as in *ba-chih-ba*), AAB (e.g., as in *ba-ba-chih*) and ABB (e.g., as in *ba-chih-chih*). Testing data included 126 (42 for each pattern) phonemically balanced trisyllabic nonwords which were created in the same way. The auditory stimuli were recorded at a sampling rate of 44.1 kHz with 16-bit sound quality and the duration of syllables was equalized to 250 ms (750 ms for each CVCVCV nonword). The input to the network was jittered cochleagrams of each auditory file. A cochleagram is a spectrotemporal representation of auditory signal designed to mimic cochlear frequency decomposition. To create a cochleagram, we first removed any surrounding silence from the audio files, and then passed each sound clip through a bank of 203 bandpass filters that were zero-phase, with varying center frequencies. Low-pass and high-pass filters were included to perfectly tile the spectrum, resulting in a total of 211 filters. The final cochleagram representation was 150 x 211 (time x frequency) (Kell et al., 2018; Feather et al., 2019). We generated the cochleagrams using Python with the numpy, scipy, and librosa libraries (Oliphant, 2007; McFee et al., 2015; Harris et al., 2020). We then created ten jittered cochleagrams for each original cochleagram by utilizing data augmentation (specifically jittering in the time domain using random sigma values between (0.03, 0.09) (Um et al., 2017). A schematic representation of the audio-to-cochleagram conversion as well as sample jittered cochleagram can be found in Fig. 2A.

**Figure 2:**
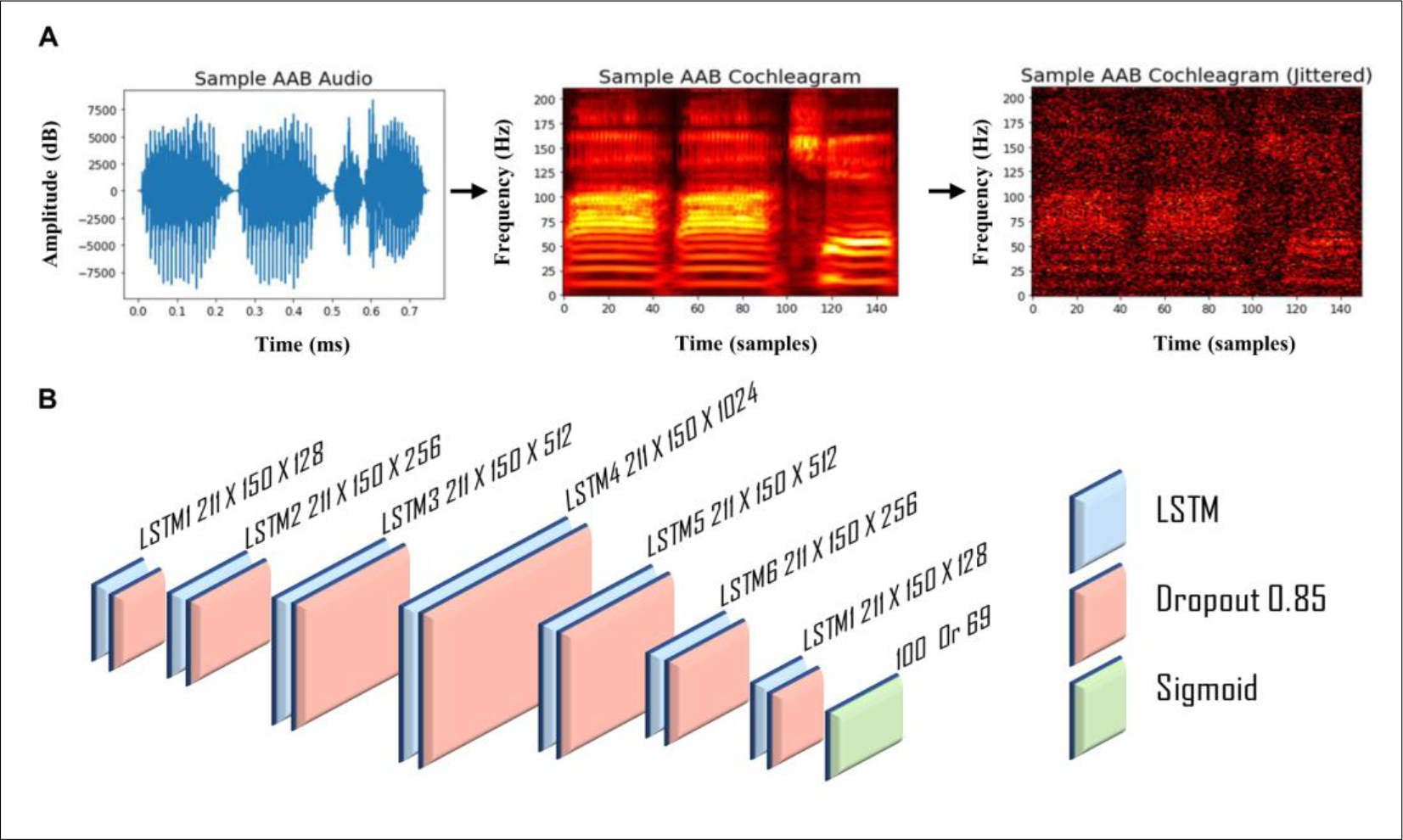
Model input and architecture. (A) Sample audio conversion to cochleagram and its jittered version. The x-axis represents the time (750 ms) and time samples (150), and the y-axis represents the amplitude (dB) and frequency (211Hz). (B) The model architecture. The model was a standard recurrent LSTM network with seven fully recurrent layers. The output layer of the model was a dense layer with the sigmoid function, either with 69 (word) or 100 (pattern) output vectors and 23 vectors for the pretrain network.

### 3.2 Training tasks and pretraining

Two separate LSTM models were created and trained independently on the same training data (7,200 tokens for 720 words). A “word learner” network was trained to differentiate between words, and a “pattern learner” network was trained to distinguish patterns. We chose the word identification task to draw attention to whole word properties with explicitly requiring sublexical segmentation into syllables. To do this, we created target vectors using a slot -based system in which there were twenty-three slots for each syllable, a total of 69 nodes (23X3). For each word, we generated a sparse target vector with 3 of 69 selected elements set to 1 (all other elements 0), representing which of the three syllables filled the twenty-three possible slots. With this task, the word learner network used whole-word syllabic properties for efficient sound to word mapping. The pattern learner network was trained to differentiate between patterns using random vectors representing the three patterns. For each of the three patterns, we generated 100-dimensional random input vectors that implicitly represented property values across dimensions. In addition, since we also checked the influence of pretraining on network performance, we trained a network on cochleagrams representing syllables using one-hot-vectors for each of the twenty-three syllables. We used cochleagrams of each syllable in the shape of 50 x 211 (time x frequency).

### 3.3 Network architecture and testing

To model variable representation in the brain, we employed LSTMs to capture the temporal structure of auditory speech data. LSTMs are a type of recurrent neural network that are capable of retaining past inputs and outputs for an extended period, making them well-suited for processing sequential data, such as time series and natural language. Based on the work of Avcu et al. (2023) and Magnuson et al. (2020), we posit that LSTMs are a superior choice for capturing long-term dependencies in auditory speech data. The pretraining model consisted of a single LSTM layer with 512 nodes and a dense layer with 23 nodes and softmax activation function. We used categorical cross-entropy as the loss function and ADAM (Adaptive Moment Estimation) (Kingma & Ba, 2014) optimization with a fixed learning rate of 0.00001. The model was trained for 5000 epochs producing very high training and validation accuracy (over 90%).

The non-pretrained word and pattern learner models consisted of seven layers with 128, 256, 512, 1024, 512, 256 and 128 LSTM nodes respectively. On top of the LSTM layers, a dense layer with vector outputs (69 for the word and 100 for the pattern learner networks). After every LSTM layer, we used a dropout layer with 0.85 (following Prickett et al. (2022)). Dropout is a regularization method that helps generalization by forcing the model to make predictions that do not overly depend on any single feature, thus encouraging robustness and preventing overfitting. See Fig. 2B for the structure of the main networks. The word and pattern learner models with pretraining consisted of the same architecture except for an additional input LSTM layer with 512 nodes with preloaded weights coming from the pretraining. The cochleagrams of size 150 x 211 were fed into the first LSTM layer. Subsequently, the output of this layer was passed onto other layers respectively. The final layer was a dense layer that transformed the input vector X to an output vector Y of length n, where n represents the number of target classes (69 or 100). We employed the sigmoid activation function for the output layer, which returns a value between 0 and 1 and is centered around 0.5. Mean squared error loss was employed to calculate the mean of squares of errors between labels and predictions, with a batch size of 100. For optimization during training, we utilized ADAM as explained above. Each of the 720 words had ten jittered tokens, and seven of these tokens were utilized for training, while three were used for validation. For the pretraining, each syllable had two hundred tokens of which 180 were used for training and 20 were used for validation. Furthermore, the word and pattern learner networks were trained for 10,000 epochs, which involved complete iterations over the training set. The training parameters, such as the learning rate, the optimization algorithm, the loss function, etc., were adopted from Avcu et al. (2023).

We calculated accuracy of the word and pattern learner networks with and without pretraining by checkpointing every 100 epochs during the training. To evaluate the distance between the predicted target vector and the true target vector, we used cosine similarity instead of a binary cross-entropy threshold value as it is more conservative and psychologically relevant (Magnuson et al., 2020; Geiger et al., 2022). We reported the average cosine similarity for all words at every 100 epochs and for both training, validation and test data. Cosine similarity between target observed patterns was calculated for trained tokens (training accuracy), reserved alternate tokens of trained syllable patterns (validation accuracy) and tokens based on syllables that were not used during training (test accuracy).

### 3.4 Decoding

We decoded the original 720 words’ activations from the best performing model iteration to check whether representations for each word would be useful for SVM to distinguish pairwise comparisons of the mean activation time courses in the three experimental conditions: ABA vs. AAB, ABA vs. ABB, and AAB vs. ABB. While the pattern learner was trained to distinguish these three patterns from each other, the word learner was trained to identify every single word. Thus, the decoding analysis shows whether the word learner grasped any useful feature to differentiate patterns while focusing on word specific features. The hidden layer activations were extracted from each LSTM layer of the models at the final time sample (150) yielding a 720 X N vectors where N is the number of hidden units in a specific LSTM layer.

We then divided the data frames into three sub data frames where each sub data frame contained pairwise comparisons, e.g., ABA vs. AAB (e.g., 480XN). Next, we standardized activations by removing the mean and scaling to unit variance using *sklearn StandardScaler* function. We then trained and tested SVMs using cross-validation (k=10) on each sub data frame. For the SVM hyperparameters, we used the *sklearn GridSearchCV* function which accepts a dictionary of different hyper-parameters. This process selected a *kernel* parameter of *poly*, a *gamma* parameter of *1*, a *C* parameter of *1e-05*, and a *tol* parameter of *1e-5*. We reported mean decoding accuracy with standard deviation for each layer of both word and pattern learner networks with and without pretraining.

### 3.5 Representational similarity analysis

Representational similarity analysis (RSA) involves assessing the correlation between decoding accuracy, determined by SVMs applied to ROI activation vectors in the brain (comprising 8 MNE measures per ROI per timepoint), and SVMs applied to activation vectors derived from each of the 7 model layers. The neural decoding accuracy data was sourced from Gow et al. (2022), where the study utilized linear SVMs to classify MNE activation timeseries within 68 distinct ROIs. It was reported that the ROIs were subdivided into eight parts, and MNE source estimates were averaged for each subdivision, accounting for trial orientation. This resulted in eight timeseries per ROI per trial, spanning from 200 ms before stimulus onset to 1000 ms after onset. Vector normalization was applied to minimize overall activation differences, and trials were down sampled to 100 Hz and bundled into sets of 10 within each condition, which were then averaged to improve signal-to-noise ratio. This process was repeated 100 times to reduce potential sampling bias. SVM classifiers were trained for each ROI and condition pair, and accuracy was assessed using a leave-one-trial-out technique. The overall accuracy on untrained trials was determined by averaging classifier performance across subjects at each timepoint yielding 1X1200 (Accuracy x Time) vectors for each of the three comparisons for each ROI. We performed preprocessing on the neural decoding accuracy vectors by narrowing our focus to the window between 100 ms and 850 ms after the word onset. This window accounts for the 100 ms delay associated with the lag between the neural signal and word onset, making the total duration still 750 ms for words. We then averaged every ten-time samples which yielded a vector of 1X75.

Model decoding accuracy data reflects the hidden layer activations associated with the 720 words from the best performing model iteration. For each model and layer, we saved hidden unit activations with size, for example, 720 × 150 × 256, where the first dimension is the number of words, the second dimension is the number of time samples, and the third dimension is the number of hidden units. We then followed the above SVM decoding steps and calculated SVM decoding accuracyat every time step for each pairwise comparison. This process yielded three vectors of size 1 × 150 (one for each pairwise comparison) for each layer of the model. We then averaged every two-time samples which yielded a vector of 1X75. SVM accuracy functions as a measure of dissimilarity, with high accuracy in two pairwise comparisons signifying a high level of dissimilarity between the compared items. We used Spearman’s rank correlation coefficient (rho), a nonparametric rank correlation measure, to assess the similarity between the decoding accuracy vector of the model and that of the brain. To enhance the reliability of our results, we employed the Monte Carlo permutation test. This simulation technique allowed us to evaluate the likelihood of obtaining the observed correlation by chance, considering the variability in our data. It offers a valuable means of verifying result robustness and gaining insight into the uncertainty associated with the correlation coefficient. The p-values associated with each correlation coefficient are based on 10,000 permutations (see Fig. 3 for a schematic representation of SVM and RSA steps).

**Figure 3:**
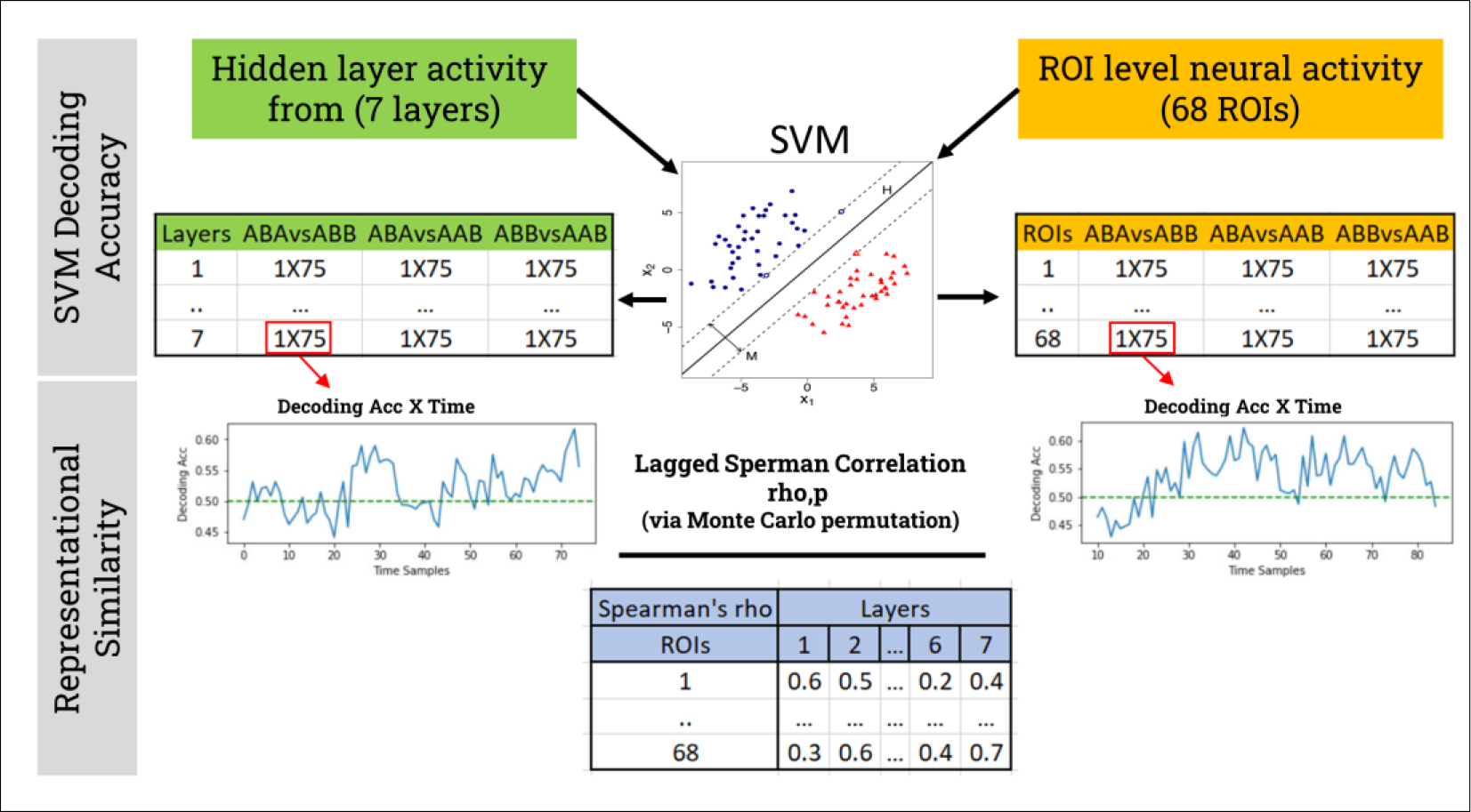
Schematic representation of SVM and RSA steps. Hidden layer activity from each layer of a specific model and ROI level neural activity from all of the 68 ROIs were fed into the SVM which outputs a decoding accuracy by time matrix for each of the pairwise comparisons. These 1X75 vectors were then correlated between the model and brain to get correlation coefficients and its associated p values. Final correlation matrix between the models and brain is created by averaging the Spearman’s *rhos* across the three pairwise comparisons.

Upon completing this procedure, we generated a matrix of dimensions 68×21 for each model, which contained correlation coefficients for every pairwise comparison across each layer (3×7). For visualization purposes, we aggregated decoding accuracy across pairwise comparisons by calculating the average of the rho values, transforming the 68×21 matrix into a 68×7 format. Since p-values cannot be averaged, we adopt a criterion where we classify a layer as “non-significant” if any p-value for a pairwise comparison within that layer exceeds 0.05. For instance, in layer 1, if the p-values are as follows: 1vs2=0.001, 1vs3=0.06, 1vs2=0.0001, we would consider layer 1 as non-significant due to the second comparison (1vs3) having a p-value of 0.06. Subsequently, we reconstructed a p-value table, designating insignificant layers with 0.1 and significant ones with 0.01. This new p-table was used for masking the insignificant correlations in the RSA plots. Finally, to compare the mean correlation values of decoding vs. non-decoding ROIs across the seven layers of each model, we used Welch’s t-test (the unequal variances t-test).

### 3.6 Replicability, hardware, and software

To confirm replicability, we repeated the entire training process for all models (including pretrained model) on separate occasions, yielding only negligible variations across iterations. Our simulations were executed on a Linux workstation equipped with an Intel(R) Xeon(R) Gold 5218 CPU operating at 2.30 GHz, supported by 98 GB of RAM, and powered by an NVIDIA Quadro RTX 8000 graphics card with 48 GB of memory. We conducted these simulations using Python 3.6, TensorFlow 2.2.0, and Keras 2.4.3. Each model required approximately 72 hours to train on this workstation, with the exception of the pretrained network, which took 6 hours.

## 4 Results

In this section, we present the outcomes of each model’s performance with and without pretraining, along with the results of SVM pattern decoding and similarity analyses in comparison to brain data.

### 4.1 Pretraining

Our premise was that seven-month-old infants are already acquainted with their language’s syllables. To assess the impact of prior knowledge on the generalization abilities of the networks, we conducted pretraining on a basic network using the twenty-three syllables employed in pattern/word learning. The outcomes of this pretraining revealed that a simple LSTM model successfully recognized all twenty-three syllables, achieving a training accuracy of 99% and a validation accuracy of 93%. This underscores that the pretrained weights, which were subsequently used for word or pattern learning, incorporate representations of these syllables.

### 4.2 Model accuracies

In our experimental setup, both a word learner, exposed to a corpus of 720 distinct words, and a pattern learner, designed to acquire three specific patterns, underwent training in two scenarios: one with pretrained weights and the other without. The results, as illustrated in Fig. 4, reveal significant disparities in their learning trajectories. In the absence of pretrained weights, both learners encountered challenges in achieving satisfactory performance levels over the 10,000 epochs. The pattern learner consistently maintained an average cosine similarity of around 0.34 throughout the entire training duration, encompassing training, validation, and test datasets. The word learner also remained relatively consistent, exhibiting a mean average cosine similarity of approximately 0.22 for training and validation accuracy (please note that test accuracy was not assessed for the word learner, given the uniqueness of each word). The pattern learner’s performance remained close to chance, while the word learner’s performance, although better than chance, remained suboptimal for a successful model. In stark contrast, when pretrained weights were utilized, both learners reached high-performance levels by the conclusion of the 10,000 epochs. The pattern learner, in particular, demonstrated an average cosine similarity of 0.71 for training, 0.59 for validation, and 0.56 for the test dataset. Notably, the assessment of test data accuracy is pivotal, as it reflects the model’s performance on novel data. The word learner also excelled, achieving average cosine similarities of 0.72 for training and 0.67 for validation data. These outcomes underscore the considerable impact of pretrained weights on the learning capabilities of our models.

**Figure 4:**
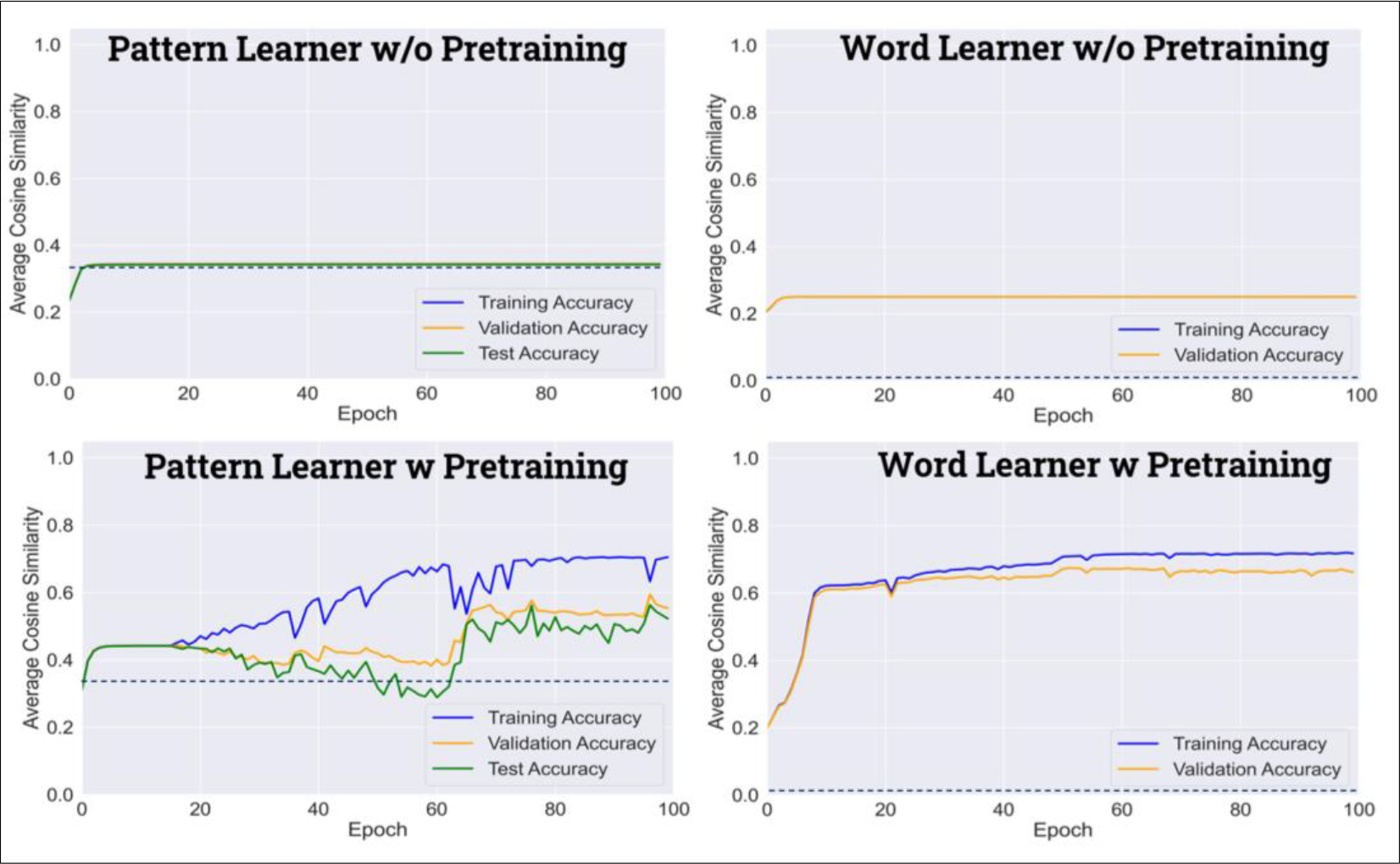
Model performance during the training of four models (word and pattern learners with and without pretraining). The top row shows the performance of models without pretraining, while the bottom row shows models with pretraining. Training performance over epochs is represented with solid lines (training accuracy in blue, validation accuracy in orange, and test accuracy in green, applicable to pattern learners only). Dashed horizontal lines indicate chance performance (33% for patterns and 0.0014% for words). The average cosine similarity between the predicted vectors and true vectors was computed for each model at every 100th epoch within the 0 to 10,000 epoch range.

### 4.3 SVM decoding accuracy

In the next phase of our experimental analysis, we employed Support Vector Machines (SVMs) to decode the hidden unit activations of both the word learner and pattern learner networks trained with and without pretrained weights. Table 1 presents the SVM mean decoding accuracy with standard deviations for each layer, focusing on the discrimination between the AAB, ABB, and ABA patterns. The results shed light on the impact of pretraining and the specific learning objectives of each model. When considering models without pretraining, we observed that both the pattern learner and word learner struggled to achieve decoding accuracy above chance levels for the AAB vs ABB comparison. This result may be attributed to the inherent repetition in both patterns. For the ABA vs AAB and ABA vs. ABB comparisons, the word learner displayed marginally better performance than the pattern learner, although both remained above chance. When considering models without pretraining, we observed that decoding accuracy varied across the layers. In particular, the pattern learner displayed increased decoding accuracy from layer 1 to layer 3, with notable improvements between layers 1 and 2. However, the performance decreased slightly in layer 4 and remained relatively consistent from layer 4 to layer 7. The word learner, on the other hand, exhibited a similar trend, with improved accuracy from layer 1 to layer 2, followed by a decrease in performance in layer 4 and consistent accuracy from layer 4 to layer 7.

**Table 1:** SVM mean decoding accuracy with standard deviation in parentheses for each layer of both word and pattern learner networks with and without pretraining. Red color reflects decoding accuracy below the chance level of 50%.

| Models | Pattern Learner w/o Pretraining |  |  | Word Learner w/o Pretraining |  |  |
| --- | --- | --- | --- | --- | --- | --- |
| Layers | ABA-AAB | ABA- ABB | AAB- ABB | ABA-AAB | ABA-ABB | AAB- ABB |
| **Layer 1:128** | 0.64 (0.04) | 0.64 (0.07) | 0.35 (0.05) | 0.79 (0.07) | 0.79 (0.04) | 0.20 (0.03) |
| **Layer 2:256** | 0.68 (0.04) | 0.69 (0.09) | 0.38 (0.04) | 0.73 (0.07) | 0.73 (0.06) | 0.24 (0.03) |
| **Layer 3:512** | 0.65 (0.07) | 0.65 (0.09) | 0.37 (0.10) | 0.77 (0.07) | 0.78 (0.06) | 0.19 (0.05) |
| **Layer 4:1024** | 0.56 (0.09) | 0.56 (0.09) | 0.45 (0.07) | 0.62 (0.10) | 0.62 (0.07) | 0.46 (0.09) |
| **Layer 5:512** | 0.56 (0.08) | 0.56 (0.08) | 0.44 (0.06) | 0.59 (0.10) | 0.57 (0.11) | 0.42 (0.07) |
| **Layer 6:256** | 0.51 (0.04) | 0.49 (0.06) | 0.40 (0.03) | 0.62 (0.07) | 0.88 (0.05) | 0.40 (0.04) |
| **Layer 7:128** | 0.51 (0.03) | 0.51 (0.03) | 0.42 (0.03) | 0.63 (0.05) | 0.62 (0.06) | 0.38 (0.05) |
| **Mean** | **0.587143** | **0.585714** | **0.401429** | **0.678571** | **0.712857** | **0.327143** |
| Models | Pattern Learner w Pretraining |  |  | Word Learner w Pretraining |  |  |
| Layers | ABA-AAB | ABA- ABB | AAB- ABB | ABA-AAB | ABA-ABB | AAB- ABB |
| **Layer 1:128** | 0.88 (0.05) | 0.88 (0.06) | 0.56 (0.06) | 0.79 (0.04) | 0.75 (0.08) | 0.71 (0.08) |
| **Layer 2:256** | 0.88 (0.03) | 0.87 (0.04) | 0.55 (0.06) | 0.83 (0.05) | 0.76 (0.05) | 0.72 (0.07) |
| **Layer 3:512** | 0.90 (0.04) | 0.88 (0.04) | 0.52 (0.06) | 0.90 (0.03) | 0.82 (0.10) | 0.68 (0.08) |
| **Layer 4:1024** | 0.97 (0.02) | 0.95 (0.02) | 0.92 (0.03) | 0.95 (0.03) | 0.95 (0.03) | 0.93 (0.05) |
| **Layer 5:512** | 0.95 (0.03) | 0.91 (0.04) | 0.93 (0.04) | 0.86 (0.04) | 0.92 (0.03) | 0.82 (0.04) |
| **Layer 6:256** | 0.95 (0.03) | 0.92 (0.04) | 0.94 (0.04) | 0.81 (0.04) | 0.88 (0.05) | 0.88 (0.03) |
| **Layer 7:128** | 0.96 (0.04) | 0.92 (0.04) | 0.94 (0.02) | 0.82 (0.05) | 0.86 (0.04) | 0.84 (0.03) |
| **Mean** | **0.927143** | **0.904286** | **0.765714** | **0.851429** | **0.848571** | **0.797143** |

In contrast, models with pretrained weights exhibited noteworthy differences. The pattern learner surpassed the word learner in the ABA vs AAB and ABA vs. ABB comparisons, displaying high decoding accuracy. In the AAB vs ABB comparison, both models achieved accuracy levels significantly above chance. Notably, the word learner demonstrated superior performance in this specific comparison compared to the pattern learner. As for the progression of decoding accuracy between layers, both the pattern and word learners demonstrated consistent high decoding accuracy across all layers, with the highest performance achieved in layer 4. These findings highlight the distinct learning dynamics of the word learner, which was primarily trained to identify individual words, and the pattern learner, designed to discriminate among the three distinct patterns. Pretraining significantly boosted the decoding accuracy of both models, underscoring the beneficial role of pretrained weights in enhancing learning capabilities. The results emphasize the importance of considering the specific objectives of neural network models and the impact of pretraining on their performance.

### 4.4 Representational similarity analysis

In addition to the decoding analysis described earlier, we conducted a comprehensive comparison of the decoding accuracy by time vectors extracted from the hidden unit activations of each layer within our models with neural activity derived from the 68 distinct ROIs. Our primary objective was to elucidate the close correspondence between human neural data and model performance in relation to pretraining and task-specific capabilities. The findings, depicted in Figs. 5 and 6, demonstrated that both the pattern and word learner models without pretraining exhibited moderate positive correlations with the neural data, particularly in the third layer of both model architectures. Notably, the regions of interest (ROIs) displaying these correlations included L-MTG_5, R-ITG_2, and R-STG_4 for the pattern learner (Fig. 5 left panel), and L-ITG_1, L-MTG_5, L-postCG_1, L-STG_1, R-ITG_2 and 3, R-MTG_2, R-STG_1, R-STG_4, and R-STS_1 for the word learner (Fig. 5 right panel). While none of the ROIs demonstrating moderate correlations with the pattern learner were decoder ROIs reported in Gow et al. (2022), it’s noteworthy that three of the ROIs showing moderate correlation with the word learner functioned as decoders, suggested to store reduplication patterns. In the case of models with pretraining, the outcomes reveal remarkably distinct patterns of correlations. Notably, the majority of decoder ROIs (with the exception of R-STG_3) and several others, demonstrated notably high correlations with the pattern learner, particularly in the later layers, while the first layer did not show any significant correlation. Conversely, for the word learner, we observed a contrasting trend, wherein all decoder ROIs and numerous additional regions exhibited substantial correlations primarily with the initial layers, while the final layer displayed comparatively weaker correlations. In addition, mean correlations between the seven layers of each model and decoder ROIs vs non-decoder ROIs (Fig. 7) showed that in all four models across all seven layers, decoder ROIs showed higher correlation than non-decoder ROIs and these correlations are significantly different from each other according to the Welch’s t-test.

**Figure 5:**
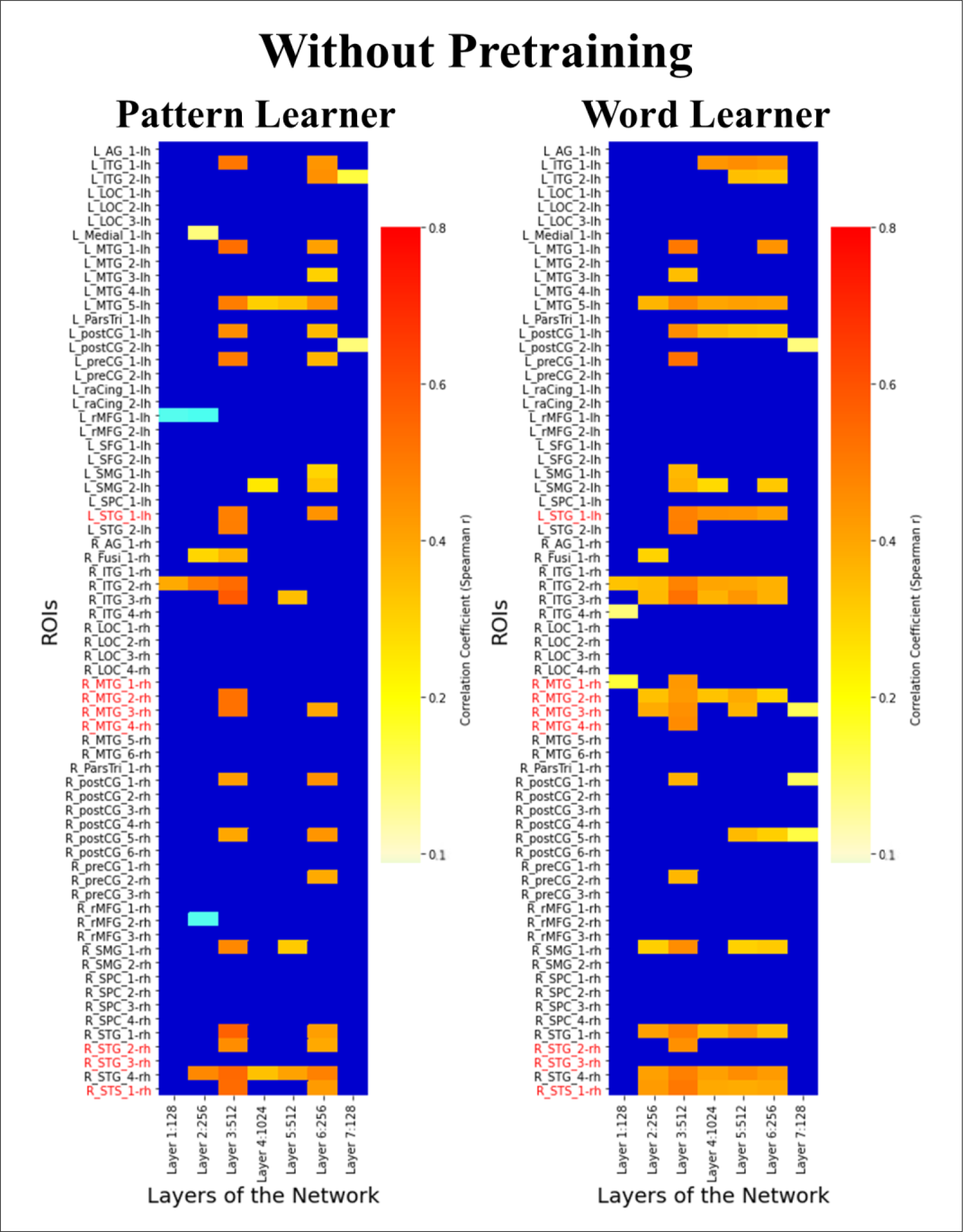
Heatmaps illustrating the correlation between SVM-based decoding accuracy applied to ROI activation vectors in the brain and SVMs applied to activation vectors across the 7 layers in the pattern and word learner models without pretraining. Each cell within the heatmap represents the correlation (Spearman’s rho) between the decoding accuracy time vector of an ROI and that of a layer in the model. Insignificant correlations are masked by blue shading. Decoder ROIs from Gow et al. (2022) are marked with red color.

**Figure 6:**
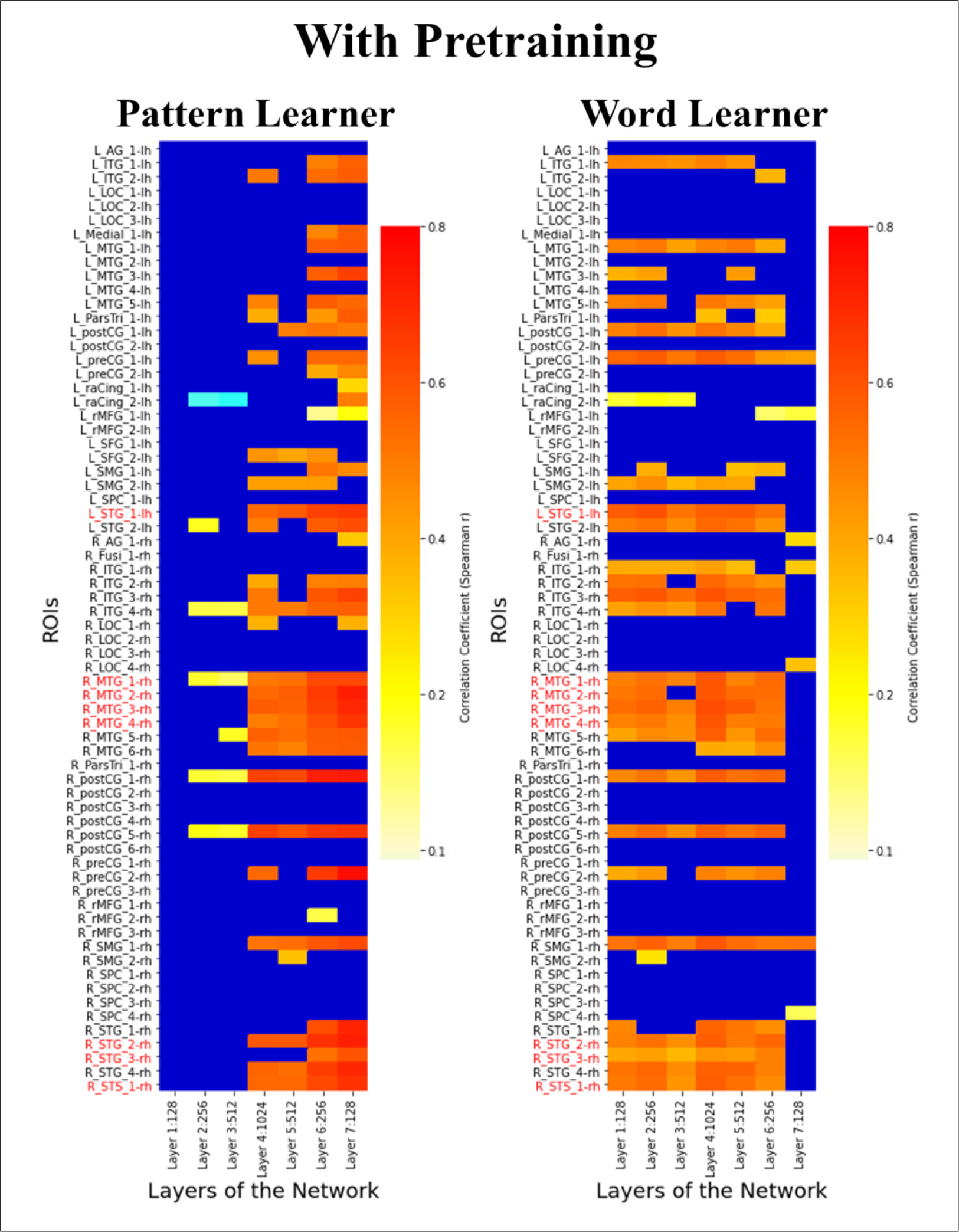
Heatmaps illustrating the correlation between SVM-based decoding accuracy applied to ROI activation vectors in the brain and SVMs applied to activation vectors across the 7 layers in the pattern and word learner models with pretraining. Each cell within the heatmap represents the correlation (Spearman’s rho) between the decoding accuracy time vector of an ROI and that of a layer in the model. Insignificant correlations are masked by blue shading. Decoder ROIs from Gow et al. (2022) are marked with red color.

**Figure 7:**
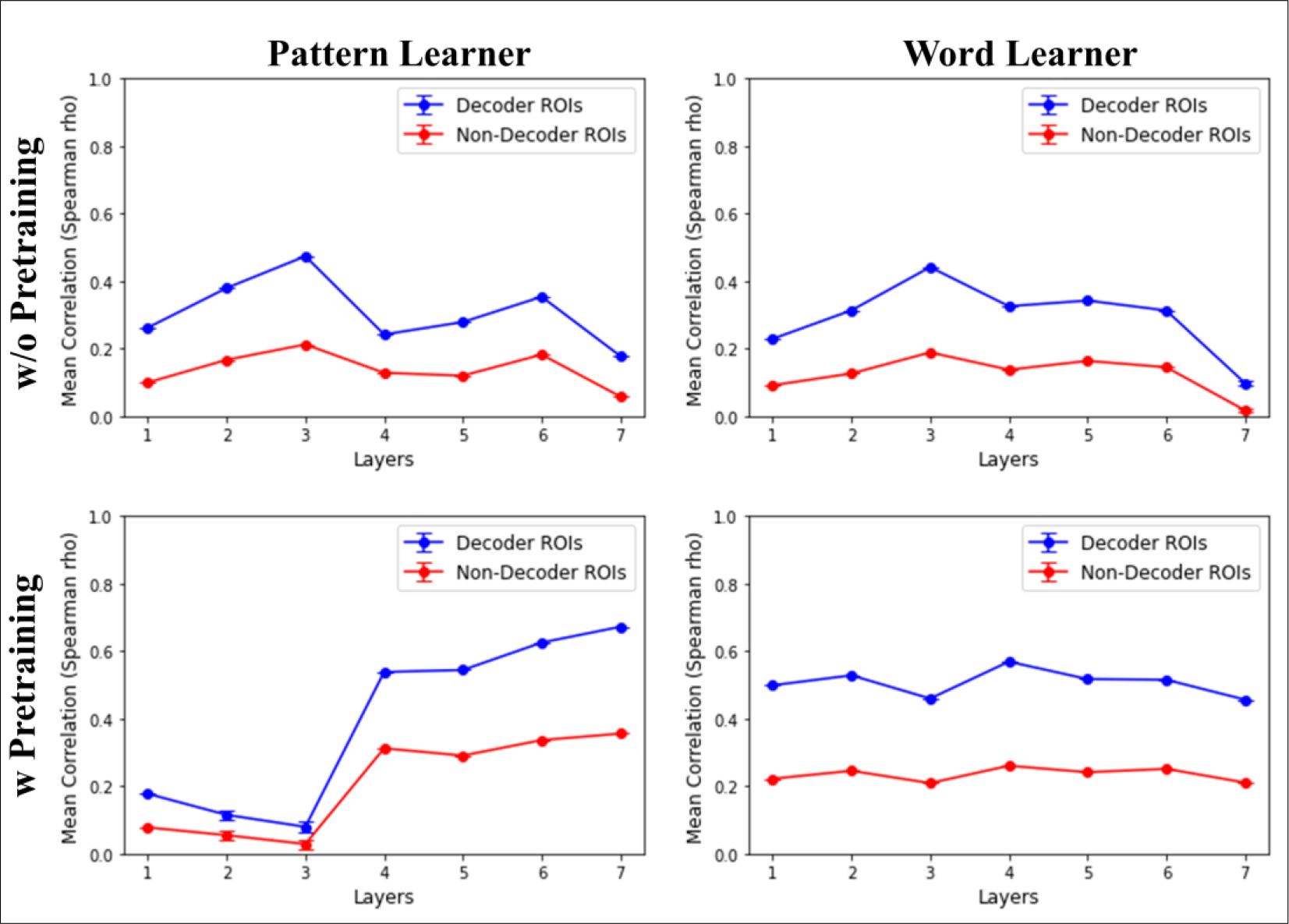
Mean correlations between the seven layers of each model and decoder ROIs vs non-decoder ROIs. Top row shows the models without pretraining, and bottom row shows the models with pretraining. Mean correlations (Spearman’s rho) for decoder ROIs are shown with blue color and non-decoder ROIs with red color. Error bars represent the Welch’s t-test p-values, which indicate the statistical significance of the mean differences of correlation between decoder and non-decoder ROIs for each layer.

## 5 Discussion

Generativity, a fundamental aspect of human language and cognition, has been the subject of an extensive investigation in both linguistic theory and computational modeling. Our study delved into this intricate aspect by employing deep associative models to investigate whether the neural abstract representations that support generativity align with the representations discovered by the variable-free model. To do this, we examined the role of pretraining and task-specific performance in mimicking cognitive generativity, particularly in the context of repetition-based rules, and drawing connections to human neural data. Specifically, we explored how task specificity and pretraining impact the performance of associative models, drawing connections between these models and human neural data obtained through MR-constrained simultaneous MEG/EEG.

Our investigation initially aimed to understand the role of pretraining in modeling generative abilities. To do this, we trained deep LSTM models both with and without pretraining, considering the premise that seven-month-old infants possess some prior knowledge about their language’s syllables. The results of our pretraining analysis underscored the substantial impact of prior knowledge, as models pretrained on syllables exhibited remarkable performance improvements, demonstrating that pretraining not only improves training accuracy but also enables models to excel on novel data. This finding resonates with prior research highlighting the influence of prior knowledge in the context of generative rule learning (Seidenberg & Elman, 1999a, b; Altmann, 2002; Geiger et al., 2022; Prickett et al., 2022) and offers valuable insights into the learning dynamics of neural network models. These insights can potentially be extended to the understanding of early language acquisition in infants.

The subsequent examination of model performance unveiled intriguing dynamics concerning the learning trajectories of word learners and pattern learners. Without pretraining, both word learners and pattern learners faced challenges in achieving reliable performance. The consistency of their average cosine similarities throughout training indicates the difficulty these models had in generalizing repetition patterns from untrained weights. These findings emphasize the complexities of repetition-based rule learning, even for models, and shed light on the intricate nature of human generativity. Moreover, the results with pretrained weights indicated that both categories of models achieved high levels of performance indicating the capacity to discern repetition patterns effectively.

Furthermore, the application of SVMs for decoding the hidden unit activations revealed critical insights into the representations of the repetition patterns within our models. Notably, models without pretraining displayed moderate positive correlations with neural data, especially within the third layer. The alignment of neural data and model performance highlights the potential of these models to capture aspects of human cognitive processing. It also underscores the importance of considering layer-specific dynamics when interpreting model representations. However, the difference between the pattern and word learner models, especially when pretrained, stood out. The pretrained pattern learner exhibited high correlations with decoder ROIs, especially in later layers, while the pretrained word learner displayed strong correlations with the initial layers. In addition, the consistent trend of decoder ROIs showing higher correlations compared to non-decoder ROIs across all layers reinforces the model’s capacity to simulate the cognitive generativity observed in human neural data.

These results lead to an intriguing question: why do pretrained word and pattern learners exhibit distinct behaviors in decoding ROIs across layers? The divergence between pretrained word and pattern learners, particularly in terms of correlations between early and later layers, may be attributed to differences in their learning objectives and strategies. The word learner, focused on individual word recognition, may prioritize early layers to capture fine-grained acoustic and phonetic features critical for word identification. In contrast, the pattern learner, tasked with recognizing abstract repetition patterns, may rely on later layers to capture more complex, higher-level representations necessary for this task. Deep neural networks often exhibit hierarchical learning, with early layers capturing low-level features and later layers capturing abstract ones, leading to varying correlations with neural data. Overfitting during training and the complex nature of neural data can also contribute to the observed differences. Further research is needed to explore the specific representations in different layers and their alignment with neural processes related to word recognition and pattern learning in the human brain.

In light of our findings, it is essential to recognize the limitations of our study. While we have drawn parallels between our models and human neural processes, these models remain simplifications of the complex neural systems of the human brain. Furthermore, our analysis was centered on a specific task related to repetition patterns. Exploring a broader range of linguistic and cognitive tasks would offer a more comprehensive understanding of the capabilities of these models. Future research could explore various aspects of generative rule learning, including the integration of multiple linguistic cues, the role of hierarchical feature representation in pretraining, and the extent to which generative models can replicate aspects of cognitive generativity. By embracing these challenges, we can continue to bridge the gap between computational models, human behavior, and the neural processes that underlie generativity in language and cognition.

In conclusion, our results suggest that associative mechanisms operating over discoverable representations capturing abstract stimulus properties account for a critical example of human cognitive generativity highlighting the crucial significance of generative AI models in simulating and understanding cognitive generativity within the realms of human learning and representation.

## Acknowledgments

We would like to thank John Rho for helping us run the early versions of the models, and Skyla Lynch for assisting in the representational similarity analysis.

## Conflicts of Interest

The authors declare that the research was conducted in the absence of any commercial or financial relationships that could be construed as a potential conflict of interest.

## Funding

This work was supported by the National Institute on Deafness and Other Communication Disorders (NIDCD) grant R01DC015455 (P.I.: Gow).

## Notes

### Competing Interest Statement

The authors have declared no competing interest.

